# The Urogenital *Lactobacillus* Axis Across Menopause: Connections Between the Vagina, Bladder, and Urinary Metabolome

**DOI:** 10.64898/2026.09.18.751586

**Authors:** Nazema Y Siddiqui, Nicole M Diaz, Lindsey Burnett, Jasmine Zemlin, Li Ma, Lisa Karstens

## Abstract

Alterations in the female urinary microbiome have been linked to several common urinary conditions that increase in prevalence after menopause. Here we use sequencing and metabolomic data to compare the urinary bladder microenvironments between women who are premenopausal, postmenopausal, and postmenopausal using vaginal estrogen. While the urinary microbiome differs between pre- and postmenopausal women, at least 6 weeks of vaginal estrogen is associated with higher urinary lactobacilli and fewer opportunistic pathogens in the urinary postmenopausal microbiome, which closer approximates what is seen in the premenopausal context. Changes in urinary *Lactobacillus* species are correlated with vaginal pH and the same *Lactobacillus* species found in the vaginal microbiome tend to also be identified within the urinary bladder microbiome. Urinary metabolites differ by context (e.g., pre-, post-, and post-menopausal using vaginal estrogen), but the majority of metabolites that drive these differences are unannotated. Of the metabolites with shared associations among different *Lactobacillus* species, *L. iners* have consistently inverse relationships compared to *L. crispatus, L. gasseri*, or *L. jensenii*.

## Introduction

The human urinary microbiome refers to microorganisms residing in the urinary tract. While several human niches have undergone detailed microbial characterization, it is only over the last decade that prior dogma suggesting sterility of the bladder was disputed, and the urinary microbiome was confirmed to exist^1,2^. We are now beginning to understand how microbial communities within this niche promote health or disease within the urinary bladder.

Recurrent urinary tract infections (UTI), urinary urgency, and urgency urinary incontinence (UUI) are bladder disorders that differentially affect females more than males, and where prevalence rises dramatically to 30-40% of women after menopause^3–8^. Sex-specific differences in urethral anatomy and hormonally driven alterations in mucosal tissue are thought to play a role in the female prevalence of these disorders. However, over the last decade, disruptions in the urinary microbiome have also been implicated. Bladder microenvironments with reduced lactobacilli and higher microbial diversity are associated with the presence of UUI, increased incontinence severity^9^, and a propensity towards infection from potentially opportunistic pathogens^10–17^. In contrast, healthy female bladder environments are associated with low biodiversity and are often dominated by lactobacilli^18,19^. Local microbial ecology and presence of lactobacilli are influenced by steroid sex hormones (e.g., estrogen) such that complex relationships are likely to exist. Indeed, age and menopause are the most significant drivers of differences in urinary microbiome composition, even among twins^20^.

In the neighboring vaginal niche, local ecology has been studied across different female life stages. Of the various *Lactobacillus* species, *L. crispatus* is most associated with health, while other species such as *L. iners* are more prevalent in altered community states and irritative conditions like bacterial vaginosis^21–24^. Species-level differences in *Lactobacillus* metabolism result in differential niche dominance, which is often based on locally available nutrient sources^25^. During the menopausal transition, the loss of estrogen triggers dramatic ecological change in vaginal microbiota^26–28^, with a precipitous drop in *Lactobacillus* genera. Given that the female vaginal and urinary microenvironments are adjacent and interconnected,^29,30^ it is not surprising that investigators have shown a significant decrease in urinary bladder lactobacilli after menopause^31^.

In contrast to the vaginal microenvironment, much less is known about the behavior of lactobacilli within the female urinary bladder. This is particularly important for human health given efforts to modulate the urinary microbiome to mitigate diseases like postmenopausal recurrent UTI. In the vaginal microbiome, the use of exogenous estrogen hormone after menopause restores the presence of *Lactobacillus* species^26–28^. However, the effects of estrogen hormones on the urinary microbiome are reportedly variable. In two studies, no changes in relative abundance of urinary lactobacilli were demonstrated after 3-4 months of vaginal estrogen therapy^32,33^. However, these studies were limited by small sample sizes with fewer than 20 participants allocated to estrogen therapy. Furthermore, these studies used vaginal tablet or ring delivery systems for local estrogen therapy, which deliver low dosages of hormone to the upper vaginal canal. In contrast, a study by Thomas-White *et al*.^34^ of 62 women using vaginal estrogen cream, which also affects the lower vagina and peri-urethral tissues, demonstrated increased urinary bladder *Lactobacillus* after 3 months of therapy^34^. A report by Neugent *et al*. corroborated these findings by showing strong associations between urinary *Lactobacillus* and estrogen hormone therapy. In contrast to Thomas-White *et al*.^34^, the report by Neugent *et al*.^14^ assessed voided urine samples where there is some obligate vulvovaginal contamination when the urine sample is collected. However, they noted a substantial increase in *Lactobacillus* when postmenopausal women used systemic estrogens (e.g., oral or patch delivery systems), and more variable increases in urinary *Lactobacillus* for those using vaginal hormone therapy^14^. Since clinicians often prescribe vaginal estrogen therapy for treatment of recurrent UTIs, clarifying whether and how hormones affect the urinary microbiome has specific relevance for clinical care.

Taken together, the current literature suggests that estrogen hormone therapy likely affects the urinary bladder by increasing the relative abundance of bladder lactobacilli, and thus decreasing the relative abundance of other bacteria, including potential pathogens, within the resident microbiome. Application of hormone at higher dosages or closer to the urethral tissue appears to matter, but the effects on the bladder microbiome need to be verified in samples without vulvovaginal contamination. To conclusively test this theory, we studied a cohort of women where urine samples were obtained directly from the bladder via catheterization under a research protocol. We specifically studied women using local vaginal estrogen cream to evaluate effects on the urinary microenvironment. We compared the bladder microbiome among premenopausal (Pre), postmenopausal (Post), and postmenopausal women using vaginal estrogen therapy (Post+VE) to identify ecologic differences across the menopausal/hormonal continuum. We used strict inclusion criteria to ensure lack of active infection and specifically excluded women with recurrent UTI since repeated infections and antibiotics leave lasting impacts on the urinary microbiome that are independent from menopausal and hormonal effects alone^14,35^. Prior metagenomic studies suggest that bacteria within the vaginal and bladder environments are more closely related than either environment is with gut^29^. Thus, we co-sampled the urinary bladder and vaginal environments to gain deeper understanding of spatial relationships in the paired ecological niches. Finally, we compared the urinary metabolites from environments dominated by various *Lactobacillus* species.

Here, we show that urinary bladder lactobacilli are reduced in relative abundance among post-compared to premenopausal women but recover after at least 6 weeks of local vaginal estrogen therapy. In contrast to the vaginal microenvironment where *L. iners* is associated with dysbiotic states, in this population *L. crispatus, L. iners*, and other *Lactobacillus* species were associated with bladder health and absence of UTI. We present novel data establishing tight correlations in the *Lactobacillus* species from paired bladder and vaginal environments and demonstrate differences in urinary metabolites that exist based on the keystone species identified within the urinary niche. Taken together, these findings corroborate the hypothesis that estrogen alters the microbiome within the urinary bladder, perhaps by increasing the relative abundance of whichever *Lactobacillus* species has baseline niche dominance within a woman’s interconnected vaginal microbiome. Different *Lactobacillus* species are associated with distinct urinary metabolites and while many *Lactobacillus* species were associated with urinary tract health in this population, it remains to be seen if certain species are more helpful for mitigating recurrent UTIs once they have begun. Future studies that gain a deeper understanding of these metabolic features may allow investigators to discern how to further enhance and support growth of different *Lactobacillus* species within the urinary tract. Ultimately, these efforts could have substantial effects in mitigating the highly prevalent problem of recurrent UTIs in postmenopausal women.

## Results

### In both the vagina and urinary bladder, microbial compositions differ between pre- and postmenopausal women but are partially restored in the presence of vaginal estrogen

After confirming inclusion and exclusion criteria (see Methods and Supplemental Figure 1), a total of 166 female participants provided catheterized urine samples, which were processed for 16S rRNA V4 amplicon sequencing (Supplemental Table 1). After removing samples that did not generate sequencing data or were filtered due to low number of sequencing reads (i.e., less than 2,000), we analyzed the urinary bladder microbiome from 149 participants, whose clinical and demographic characteristics are summarized in **Table 1**. These include 28 Premenopausal (Pre), 61 Postmenopausal (Post), and 60 Postmenopausal women using locally acting vaginal estrogen therapy for at least 6 weeks (Post+VE). Study participants also provided vaginal swab samples; 13 of these did not meet sequencing thresholds (see Methods), resulting in vaginal microbiome data for 136 participants. We used non-metric multidimensional scaling (NMDS) to visualize and compare urinary and vaginal microbial communities, estimated by weighted UniFrac distances, at the species level (**Figure 1**). Within the urinary bladder and vaginal communities, Pre and Post samples cluster separately, while Post+VE samples span across these clusters. These findings are confirmed by permutational multivariate analysis of variance (PERMANOVA) analyses that show significant differences in Post vs Pre but not Post+VE vs Pre microbiota for both urinary and vaginal niches. Since there are several clinical or demographic differences between groups that could theoretically influence microbial composition and confound results, we also added several covariates of interest (**Supplemental Table 2**) and report adjusted PERMANOVA analyses. In the bladder, when adjusting for potentially confounding variables, the microbial composition of Post women remains significantly different compared to the other groups (p=0.001) with no additional differences in microbial composition between Post+VE and Pre groups (p=0.478). Notably, in this analysis age does not demonstrate an additional association with urinary bladder microbial composition after adjusting for menopausal & hormonal status. In the vagina, the microbial composition of Post women also remains significantly different compared to the other groups (p=0.002), though age and sexual activity are independently associated with microbial composition aside from what is observed with menopause/hormonal status.

**Table 1.** Summary of Demographic & Clinical Data.

|  | ***Pre***<br>***(n=28)*** | ***Post***<br>***(n=61)*** | ***Post+VE***<br>***(n=60)*** | ***p-value*** |
| --- | --- | --- | --- | --- |
| **Study Site** |  |  |  |  |
| Duke | 24 (85.7) | 56 (91.8) | 54 (90.0) | 0.678 |
| OHSU | 4 (14.3) | 5 (8.2) | 6 (10.0) |  |
| **Age in years** | 38.5 ± 4.8 | 66.2 ± 7.1 | 66.7 ± 6.7 | **<0.001\*** |
| **Race** |  |  |  |  |
| Caucasian | 23 (82.1) | 44 (72.1) | 54 (90.0) | **0.013** |
| African American | 1 (3.6) | 13 (21.3) | 5 (8.3) |  |
| Other<sup>#</sup> | 4 (14.3) | 4 (6.6) | 1 (1.7) |  |
| **Diabetes** | 3 (10.7) | 7 (11.5) | 3 (5.0) | 0.407 |
| **BMI (kg/m<sup>2</sup>)** | 28.4 ± 6.9 | 29.0 ± 7.9 | 27.5 ± 4.3 | 0.448* |
| **Sexually Active** | 21 (75.0) | 25 (41.7) | 32 (55.2) | **0.020** |
| **Cigarette smoker** |  |  |  |  |
| Current | 7 (25.0) | 25 (41.0) | 18 (30.0) | 0.386 |
| Past | 1 (3.6) | 2 (3.3) | 0 (0.0) |  |
| **Overactive Bladder** | 7 (25.0) | 15 (24.6) | 25 (41.7) | 0.100 |
| **Daily probiotics/yogurt** | 14 (50.0) | 36 (60.0) | 28 (46.7) | 0.345 |
| **Vaginal pH<sup>^</sup>** | 4.72 ± 0.63 | 5.29 ± 0.54 | 5.16 ± 0.55 | **<0.001\*** |

Data are summarized as n(%), mean ± standard deviation, or median [interquartile range]
P values by \*one-way ANOVA or Chi-square
<sup>#</sup>Other race category represents participants who self-identified as either Asian, Native American, or Multi-racial. There were no self-identified participants of Latina ethnicity.
<sup>^</sup>Vaginal pH collected in n=28 Pre, n=57 Post, and n=34 Post+VE participants

**Figure 1.**
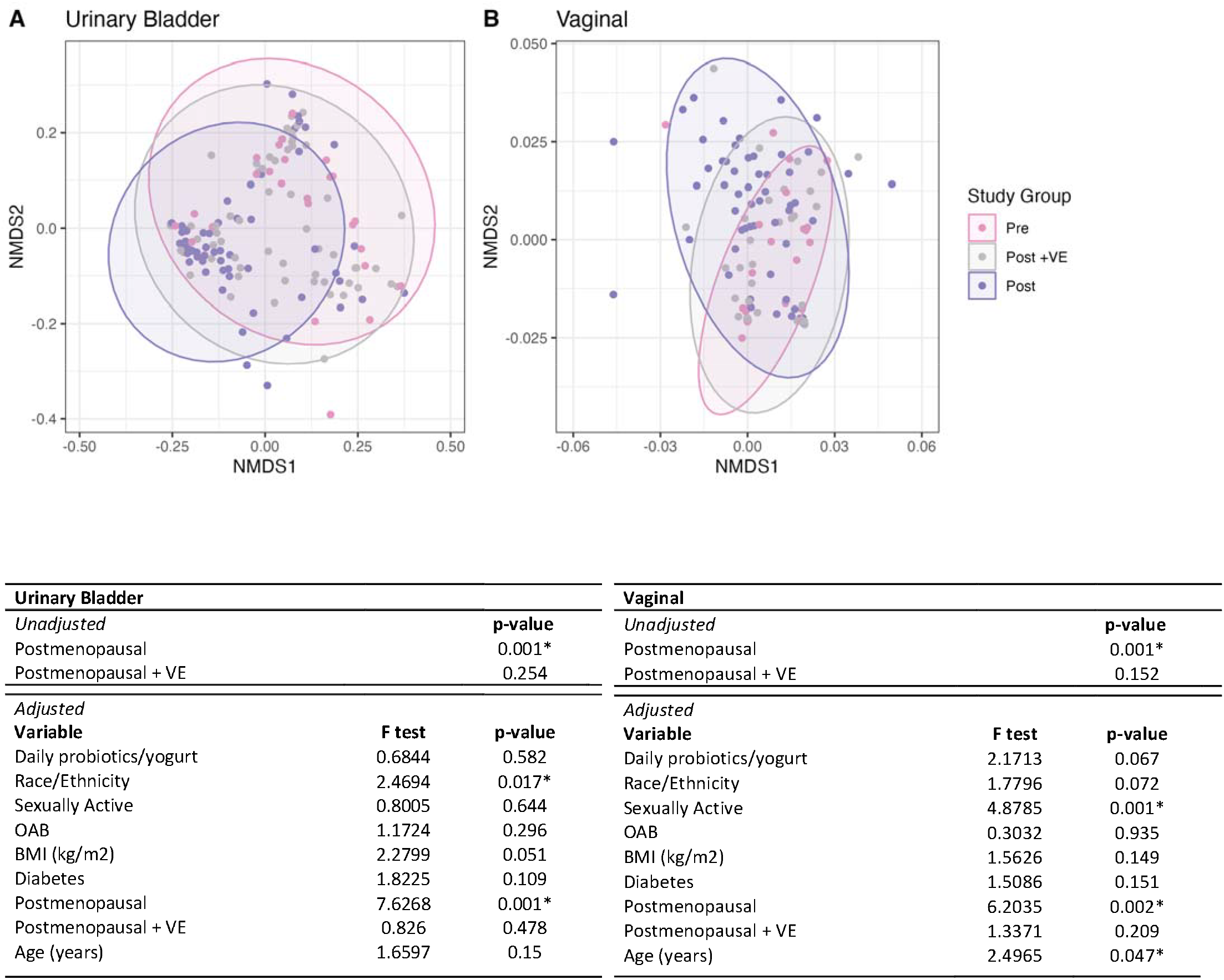
Microbial communities in. A) urinary bladder and B) vagina. Non-metric multi-dimensional scaling (NMDS) analyses comparing weighted UniFrac distances by Pre, Post, Post+VE groups at the species level. Points that are close in proximity to each other represent similar microbial communities; points further apart represent more dissimilarity. In both panel A (urinary bladder) and panel B (vagina), the Pre and Post participant’s communities cluster somewhat differently, with Post+VE spanning between the two. NMDS are unadjusted; thus permutational multivariate analysis of variance (PERMANOVA) models were constructed to compare weighted UniFrac distances by group while controlling for potential confounders. In both panels, premenopausal = reference group. Since order of variables affects measured effect, the order was specified as above and age was intentionally added after menopausal group to identify additive age-related effects over the thresholding of menopause.

### In the urinary bladder, *Lactobacillus* drives differences seen in the urinary microbiome after menopause; these differences are mediated by vaginal estrogen and vaginal pH

Within the urinary bladder, the microbiome of premenopausal women is often dominated by lactobacilli and Actinobacteria, while in postmenopausal women there is a shift away from these bacteria and towards environments dominated by Proteobacteria (**Figure 2A**). In the presence of at least 6 weeks of vaginal estrogen, the microbiome shifts back towards one more heavily dominated by lactobacilli and Actinobacteria, similar to what is seen in premenopausal women. In our analysis, *Lactobacillus* was the most frequently identified taxon within the urinary bladder, and this keystone genus appears to be a major driver of differences seen with menopause. When evaluating the median relative abundance of *Lactobacillus* (inclusive of all species), Pre is significantly higher than Post (p=0.0003), and Post is significantly lower than Post+VE (p=0.004) groups. There are no significant differences in median abundance of *Lactobacillus* between Pre and Post+VE (p>0.05). This finding corroborates the conceptual model that lactobacilli are higher in the premenopausal state, decrease after menopause, but then can recover or return to a higher abundance <u>within the urinary bladder</u> after at least 6 weeks of vaginal estrogen usage (see **Figure 2B**). Estrogen acidifies the vaginal environment and lowers vaginal pH^36^. In our study population, regardless of menopausal status, we consistently identified that lower vaginal pH was inversely correlated with higher relative abundance of *Lactobacillus* within the urinary bladder, and this relationship was more consistent than any correlations between age and urinary *Lactobacillus* (**Fig 2C and 2D**). These data lead to the conclusion that estrogen, whether endogenous in Pre women, or exogenous in Post+VE women leads to a more acidic vaginal environment that supports the presence of *Lactobacillus*, even within the urinary bladder. However, should the vaginal environment remain acidic due to other factors, relative abundance of bladder *Lactobacillus* may still be preserved.

**Figure 2.**
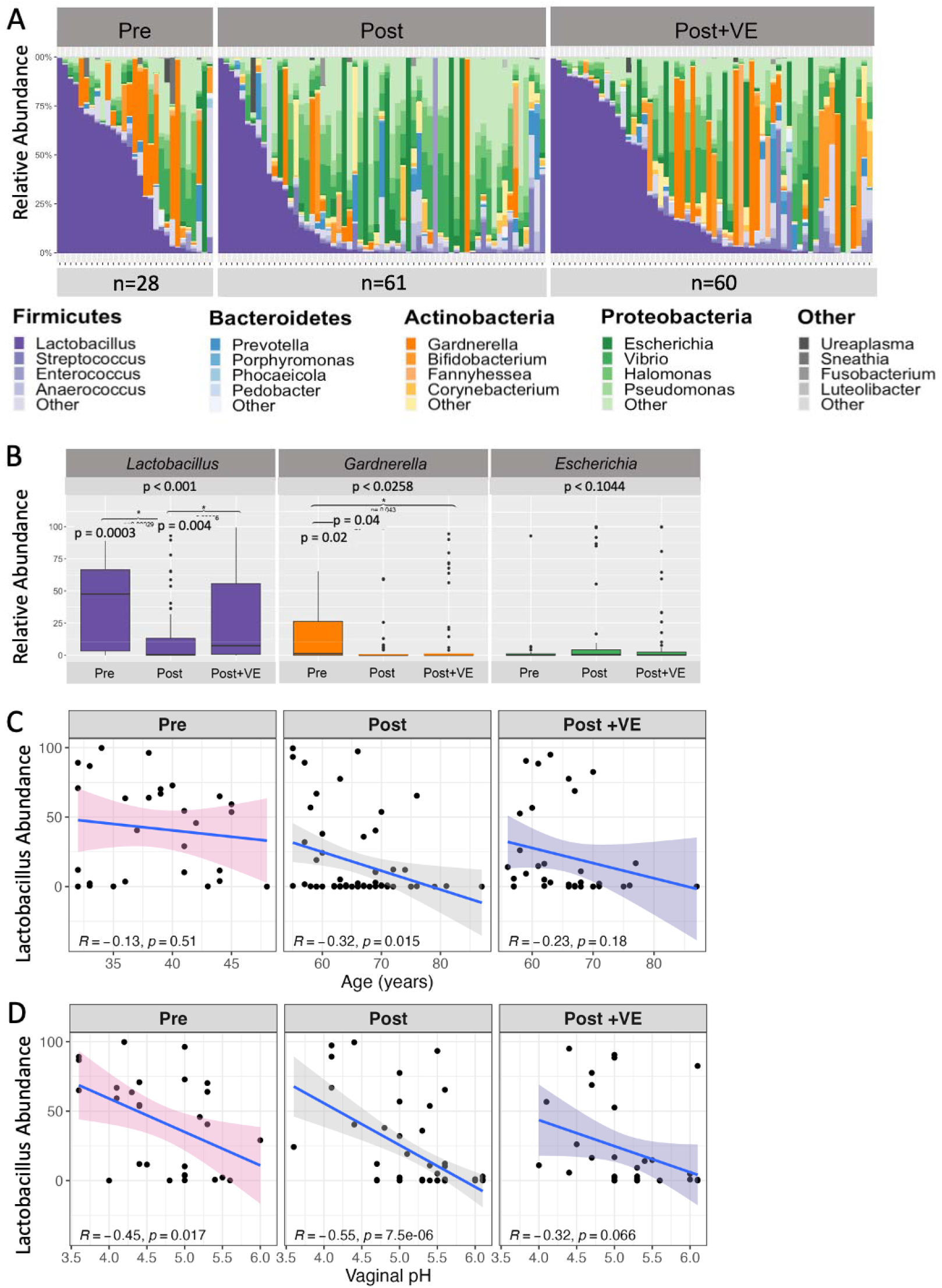
Urinary microbiome composition of female participants. A) Stacked bar plots depicting the relative abundance of microbiota (y) for each participant’s urinary microbiome (x) at the genus level, grouped by Premenopausal (Pre), Postmenopausal (Post), and Postmenopausal with vaginal estrogen (Post+VE) status. Colors depict a taxonomic Phylum with shades of a color indicating different Genera. B) Median relative abundances of key bacterial genera of interest. Relative abundance of *Lactobacillus* is significantly lower in in Post women when compared to Pre (p = 0.0003) and Post + VE (p = 0.004) groups. There are no significant differences in relative abundance between Pre and Post +VE groups (p >0.05). Median abundance of *Gardnerella* is significantly lower in both Post and Post+VE groups when compared to Pre women (p = 0.02, p = 0.04, respectively), with no significant differences between the two postmenopausal groups (Post vs. Post +VE, p > 0.05). There were no significant differences in the relative abundance of *Escherichia* between groups though abundances were low overall. C) Correlation between between age in years (x) and relative abundance of *Lactobacillus* (y) in Pre, Post, and Post+VE groups. While increasing age is negatively correlated with urinary *Lactobacilli*, this is only significant in the Post group. D) Correlation between vaginal pH (x) and relative abundance of urinary *Lactobacillus* (y) in the same groups. All three groups show a moderate negative correlation with increasing pH correlated with decreased abundance of urinary *Lactobacillus*. R denotes Pearson correlation coefficients for each plot.

### In postmenopausal women, as *Lactobacillus* decreases, potentially pathogenic genera increase within the urinary bladder microbiome

After mapping the urinary and vaginal samples to each other, we identified paired urinary and vaginal microbiome data for 124 participants. The top ten genera recovered per group from paired urinary bladder and vaginal samples are displayed in **Supplemental Table 3**. In the vagina, it is evident that the microbiome is often dominated by one genus, *Lactobacillus*. While *Lactobacillus* is the top genus in the vaginal microbiome for each phenotypic group, the median relative abundance was substantially higher in Pre and Post+VE groups compared to Post women (Median 88.3, IQR [44.94, 98.90] vs. 73.1 [7.85, 98.49] vs. 2.0 [0.10, 71.69], respectively, p=0.003). Furthermore, there was more diversity in the Post group with several additional genera identified, even in low abundance. This pattern leads to the conclusion that in the vagina, as lactobacilli decrease, other genera increase in relative abundance. While the urinary bladder microbiome appears to be more diverse and heterogeneous than the vaginal microbiome, similar phenomena are noted with lactobacilli. In the bladder, the median relative abundance of *Lactobacillus* is highest in Pre, lower in Post+VE and lowest in Post groups (Median 42.93, IQR [4.03, 66.71] vs. 7.53 [0.24, 32.75] vs. 0.55 [0.00, 14.75], respectively, p<0.001). In the Pre and Post+VE group, *Lactobacillus* remains the most abundant genus, with several potentially opportunistic bacteria present in lower abundances. In contrast, in Post women, *Lactobacillus* is no longer the taxon found in highest abundance, ranking the 6^th^ most abundant in our analysis, with nearly all other taxa in the top ten comprised of potentially opportunistic pathogens

### As *Lactobacillus* decreases, there is increased diversity within the vaginal and urinary bladder microbiomes

In contrast to the gut, where a diverse microbiome is favorable, in the vagina and urinary bladder, the premenopausal microbiome is typically low in diversity and dominated by *Lactobacillus*^26,27,31^. In instances when *Lactobacillus* decreases, other genera increase, leading to a more diverse microbiome as is seen in Post women in this study. Alpha diversity was assessed using several indices for each niche and was compared between Pre, Post, and Post+VE groups. In the urinary bladder, Post women had slightly higher alpha diversity then either Pre or Post+VE groups, though these differences did not consistently reach statistical significance, particularly when adjusting for multiple pairwise comparisons (**Figure 3**). In contrast, in the vagina, where differences in diversity indices were more pronounced, the Post microbiome was significantly more diverse, regardless of which index was used for measurement

### Vaginal and urinary bladder lactobacilli are correlated; the same *Lactobacillus* species recovered in the vagina is often found in the urinary bladder

Paired urinary bladder and vaginal microbiome data were available for 124 participants. Lactobacilli and Actinomycetia, including *Gardnerella* are positively correlated in the two niches while Enterobacteriaceae are not (**Figure 4, Supplemental Figures 2-4**). During bioinformatic processing, taxonomy was assigned at the species level using Bayesian Lowest Common Ancestor (BLCA), given that this technique improves species-level accuracy for taxonomic assignments.^33,34^ For each species annotation, the confidence score indicates the level of confidence that the correct species is assigned. The most abundant *Lactobacillus* species recovered from V4 16S rRNA sequencing were *L. iners, L. gallinarum, L. gasseri*, and *L. jensenii*, with taxonomic assignment confidence scores of 99.3%, 31%, 47.7%, and 61%, respectively. For *L. gallinarum*, which had the lowest confidence score, we hypothesized that these amplicon sequence variants (ASVs) might represent a shared sequence matching several *Lactobacillus* species. Thus, we performed Basic Local Alignment Search Tool (BLAST) searches with the ASV sequences that were called as *L. gallinarum* by BLCA. We confirmed that these ASV sequences matched to three *Lactobacillus* species (*L. crispatus, L. acidophilus, L. gallinarum*) equally, since the V4 region of the 16S rRNA gene for these three species are nearly identical. Thus, with V4 amplicon-based sequencing, we are unable to distinguish whether our study population had *L. gallinarum* or another species with an identical V4 region (e.g., *L. crispatus* or *L. acidophilus*). Since the species of *Lactobacillus* may have implications for metabolism and function, we elected to re-sequence a subset of n=71 samples using LoopSeq, which is a synthetic long read technology that provides information on the full length 16S rRNA gene^37^. With long-read data, we expected to classify lactobacilli with enhanced accuracy and precision in both urinary and vaginal samples. As we had suspected, with this technique we confirmed that what was assigned as *L. gallinarum* in the V4 amplicon-based analysis was actually *L. crispatus*. We confirmed the species recovered in the urinary bladder (see **Supplemental Table 4**) and were able to further assess the *Lactobacillus* species in matched urinary bladder and vaginal samples. When aligning *Lactobacillus* species from the urinary bladder against the vaginal data from the same participant, an obvious pattern emerges where the same *Lactobacillus* species are found in the two neighboring niches (**Figure 4**)

### Urinary metabolites differ based on the dominant *Lactobacillus* species identified in the bladder

To further investigate how the bladder environment may interact with presence of *Lactobacillus*, we performed untargeted metabolomic analyses on n=154 urine samples from 31 Pre, 67 Post, and 56 Post+VE participants. In total, 2,487 metabolite features had a Variable Importance in Projection (VIP) score > 1. Most were unannotated, with only 180 (~7%) matched to library spectra through the Global Natural Products Social 2 (GNPS2) feature-based molecular networking (FBMN) tool. Principal component analysis (PCA) and partial least-squares discriminant analysis (PLS-DA) of urinary metabolite profiles showed modest separation by study group (**Figure 5**). This separation was statistically supported by PERMANOVA, which indicated a small but significant difference in metabolite composition across groups (F (2,151) = 1.27, R^2^ = 0.017, P = 0.010). Since different *Lactobacillus* species have different nutrient and carbon sources, we hypothesized that different urinary *Lactobacillus* species would correlate with different urinary metabolites. Thus, within each group, we used the CorrOmics tool to correlate the relative abundances of the dominant *Lactobacillus* species with urinary metabolite abundances^38^. In these group-stratified analyses, we identified multiple metabolites that were positively or negatively correlated with different *Lactobacillus* species. The direction and magnitude of correlations varied by clinical group (e.g., Pre-, Post-, and Post+VE), and the vast majority of metabolic features that we identified to be correlated with lactobacilli were un-annotated (**Figure 5**). In the pre-menopausal context, where there were the highest abundances of lactobacilli, we identified several metabolites that were correlated with more than one *Lactobacillus* species. One metabolite demonstrated shared positive correlations with *L. crispatus* and *L. jensenii*. Between *L. jensenii* and *L. gasseri* there were 7 metabolites with shared correlational relationships; six metabolites were positively correlated with both species, and one metabolite was negatively correlated with both species. *L. iners* were unique in that 22 metabolites were correlated with *L. iners* and one or more other *Lactobacillus* species. In every instance, there were inverse relationships where a metabolite was either positively correlated with *L. iners* and negatively correlated with other *Lactobacillus* species, or vice versa.

## Discussion

Consistent with our hypotheses, we identified a significant reduction in *Lactobacillus* abundance in the urinary bladder microbiome of postmenopausal, when compared to premenopausal women. However, for postmenopausal women using vaginal estrogen for at least 6 weeks, abundances of urinary lactobacilli were not significantly different than the premenopausal state. These data suggest that vaginal estrogen is associated with a preservation of lactobacilli, not only in the vaginal niche as has been previously demonstrated, but in the urinary bladder as well. There appears to be a large thresholding effect such that menopausal and hormone status, perhaps more than general aging, greatly influences microbial community structure in both the vagina and female urinary bladder. In postmenopausal women, lactobacilli decrease substantially in the vagina, allowing other bacteria to increase their abundance, and thus resulting in increased vaginal microbial diversity. Here, we also present new data showing that the same phenomena occur within the female urinary bladder. In the bladder, concurrent with a decrease in lactobacilli, we see increased abundances of several potential urinary pathogens, even in the absence of clinical infection. This effect is most pronounced in postmenopausal women without estrogen therapy. However, in postmenopausal women using vaginal estrogen, there was some preservation of urinary lactobacilli, and thus reduced abundances of potentially opportunistic pathogens within the urinary microenvironment. Notably, vaginal lactobacilli are positively correlated with the presence of urinary bladder lactobacilli, and the same *Lactobacillus* species are identified in both microenvironments of the same participants, verifying the interconnected nature of the two neighboring niches, which has been previously suggested^29^.

**Figure 3.**
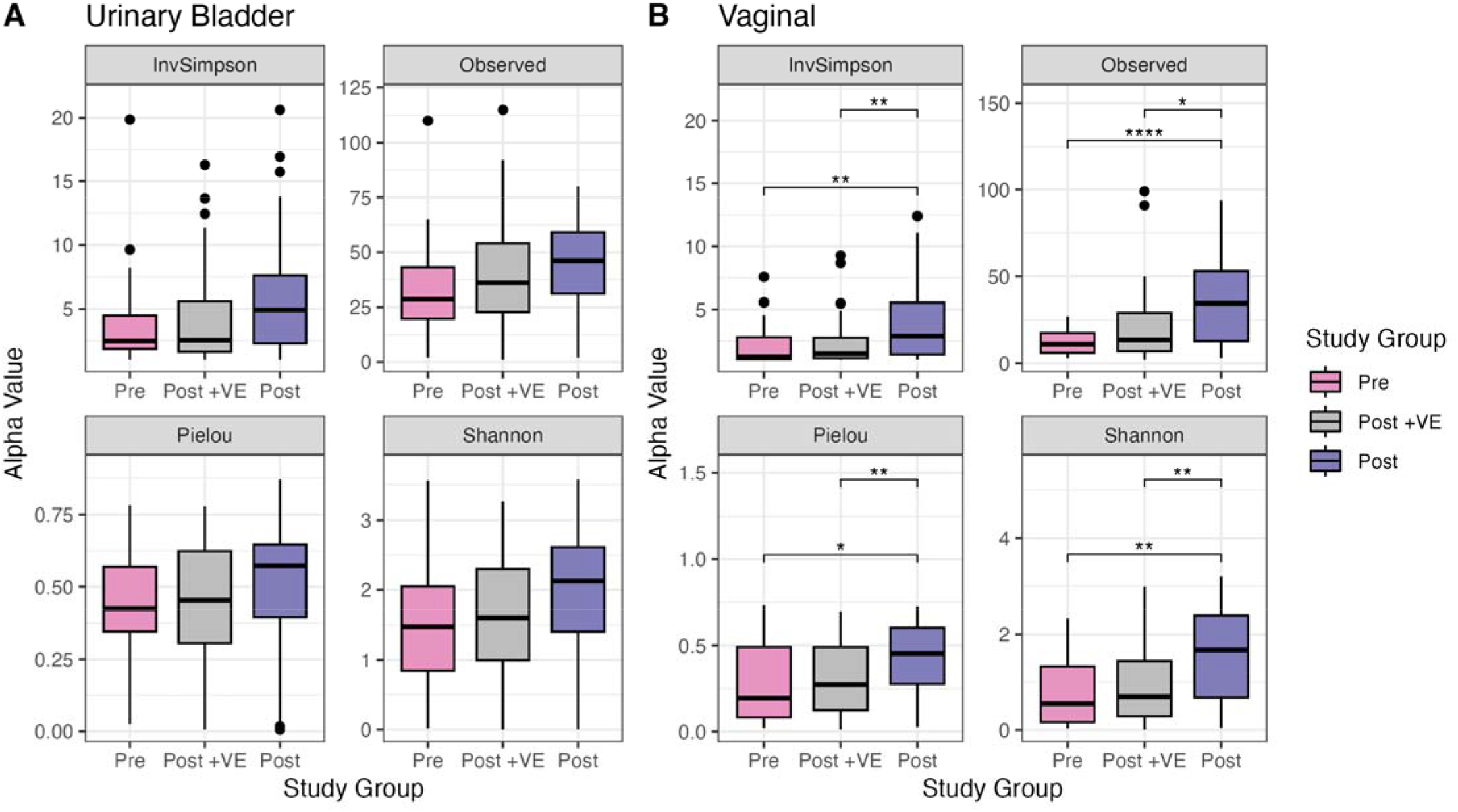
Alpha Diversity indices for Urinary Bladder and Vaginal niches. Four commonly used alpha diversity indices were calculated using the vegan,^52^ phyloseq,^53^ and microbiome^54^ R packages. These include Inverse Simpson (InvSimpson) and Shannon indices, which are comprised of richness and evenness, number of observed species (Observed), which is a measure of richness, and Pielou’s evenness (Pielou). Median diversity with interquartile ranges are summarized in plots. For all indices, median diversity is higher in postmenopausal women compared to premenopausal women or postmenopausal women using vaginal estrogen. In the urinary bladder (A), these differences did not reach statistical significance (Inverse Simpson: p= 0.04, pairwise comparisons p.adj > 0.05 | Observed: p= 0.04, pairwise comparisons p.adj > 0.05 | Pielou: p= 0.13 | Shannon: p= 0.06). In contrast, in the vagina (B), differences in diversity indices were more pronounced, and the Post microbiome was significantly more diverse, regardless of which index was used for measurement (Inverse Simpson: p = 0.001 | Observed: p= 0.002 | Pielou: p=0.003 | Shannon: p=0.004). P values calculated by Kruskal Wallis. Pairwise comparison adjusted p values noted as follows: none: p.adj >0.05; ^*^: p.adj <= 0.05; ^**^: p.adj <= 0.01; ^***^: p.adj <= 0.001;^****^: p.adj <= 0.0001.

**Figure 4.**
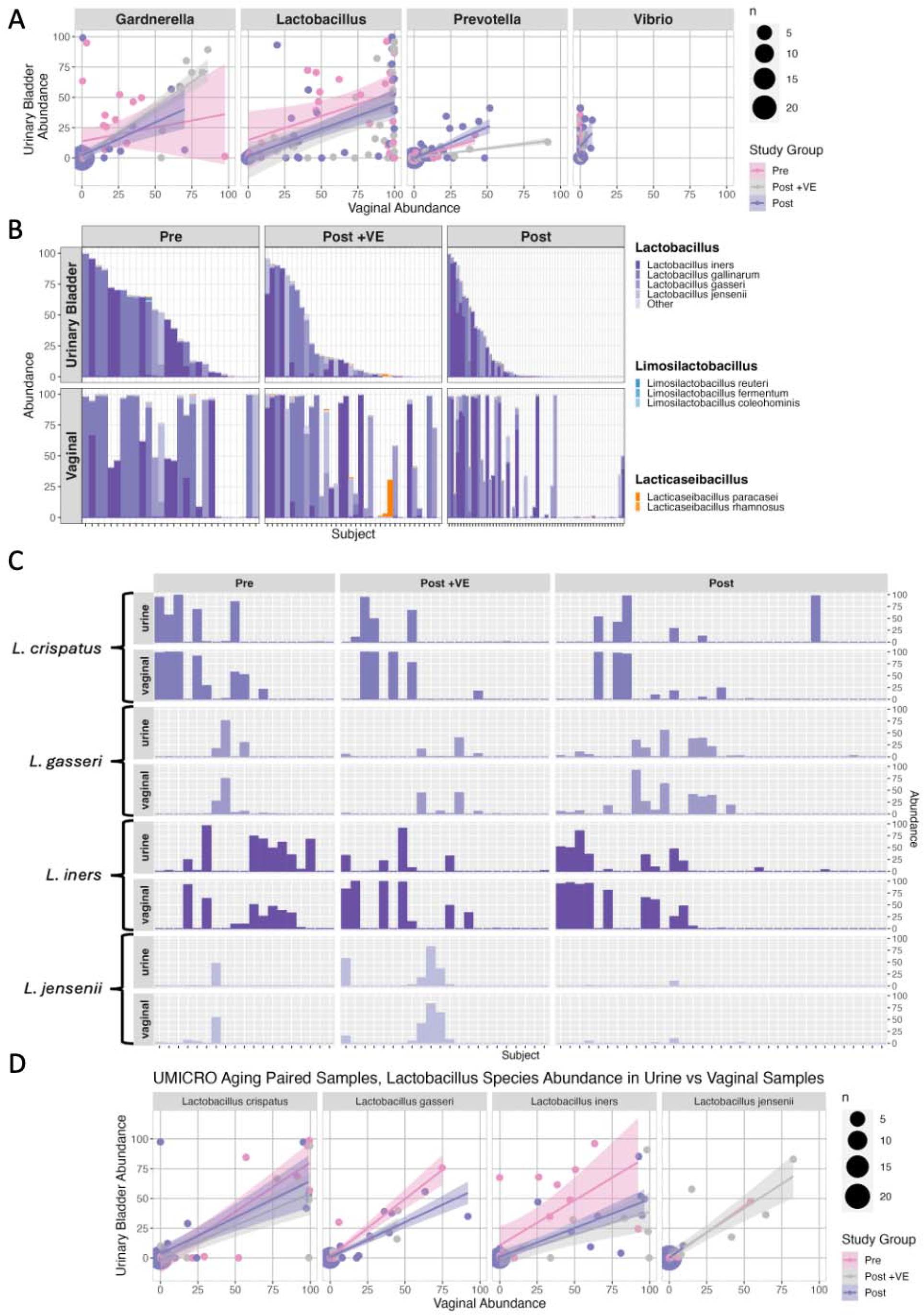
Paired relationships between urinary and vaginal microbiota. (A) Correlation plots of relative abundances of selected genera between vaginal (x) and urinary bladder (y) niches in n=124 paired samples. Data from V4 region of the 16S rRNA gene (Illumina) with taxonomy assigned using BLCA. *Gardnerella* and *Lactobacillus* demonstrate positive correlations between the adjacent niches. In contrast, *Vibrio*, a member of the phylum Proteobacteria is not correlated between niches. (B) *Lactobacillus* is further exploded to illustrate relative abundances by species in the urinary bladder (top) and aligned with the vaginal (bottom) species recovered from the same participants. Since 2020, what was once termed as *Lactobacillus* are now 24 different genera including *Limosilactobacillus, Lacticaseibacillus*, and several others.^51^ Using V4 amplicon sequencing, *L. gallinarum* is 100% identical to both *L. crispatus* and *L. acidophilus* reflecting confidence scores of 99.3%, 31%, 47.7%, and 61% for assignments to *L. iners, L. gallinarum, L. gasseri*, and *L. jensenii*, respectively. (C) Given the poor confidence of some of these species-level assignments, paired urinary bladder and vaginal samples from n=71 participants were re-sequenced using a synthetic long-read approach that covers the entire 16S rRNA gene (LoopSeq). Aligned relative abundances by confirmed species in the urinary bladder (top) and vaginal (bottom) niches of the same participants are displayed. (D) Correlation plots for each of the species in panel C showing strong positive correlations in *Lactobacillus* species between the vagina (x) and urinary bladder (y) for all phenotypic groups (Pre, Post+VE, Post) though low overall presence of *L. jensenii* limits conclusions for this species.

**Figure 5.**
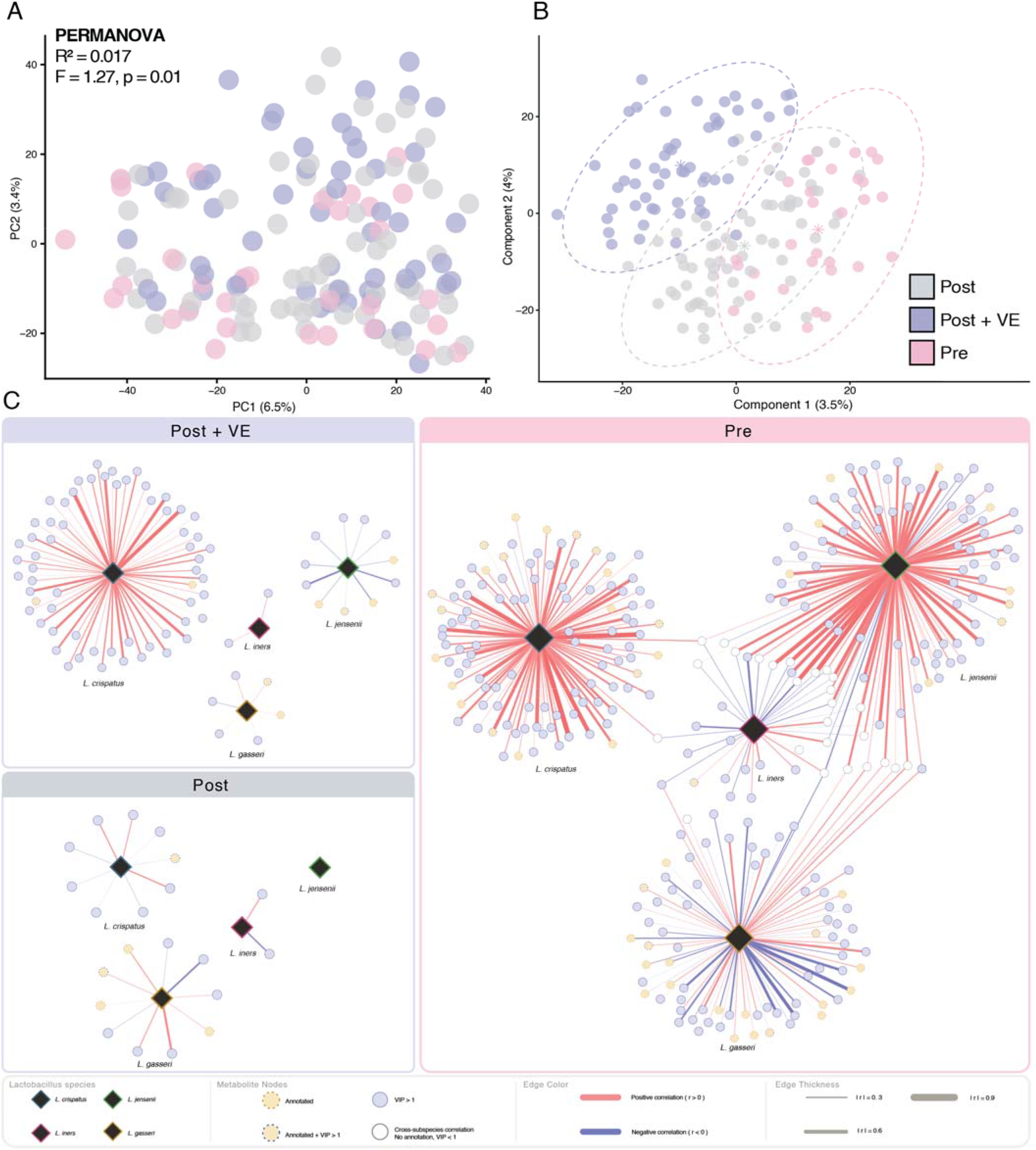
Urinary metabolite profiles and Lactobacillus–metabolite networks in Pre, Post, and Post+VE groups. Participants were grouped as Pre (premenopausal), Post (postmenopausal), and Post+VE (postmenopausal with vaginal estrogen). **(A)** Principal component analysis (PCA) of RCLR-transformed metabolomic features stratified by study group depicting overall urinary metabolite profiles (PERMANOVA, 999 permutations: *F*= 1.27, *R*^2^ = 0.017, *P* = 0.010). (**B)** Partial least-squares discriminant analysis (PLS-DA) of RCLR-transformed metabolomics features stratified by group. Dashed ellipses denote 95% confidence regions and asterisks mark group centroids. Model performance was assessed by cross-validation, showing modest but above-chance discrimination (BER = 0.56, chance = 0.67) **(C)** Correlation-based multi-omic integration (CorrOmics) performed separately within each group, linking the relative abundance of four *Lactobacillus* species (*L. crispatus, L. gasseri, L. iners*, and *L. jensenii*) to urinary metabolites (nodes). Edges represent significant *Lactobacillus* species–metabolite correlations. Networks are drawn at a common scale, so differences in size reflect the number of significant correlations per group.

Our findings are strengthened by the study design, which included intentional sampling of the bladder with catheterization to avoid vulvovaginal contamination, strict inclusion and exclusion criteria that accounted for menopausal status and vaginal estrogen, and inclusion of other potentially confounding clinical covariates in our analyses. We intentionally excluded women ages 48-54 in perimenopause, since this could add unnecessary heterogeneity to our findings. We excluded active UTI or history of recurrent UTI to ensure our data reflect a nonpathogenic urinary microbiome. Furthermore, by using paired urine and vaginal samples from the same participants, and through improvements in sequencing technologies, we provide novel detailed data about urinary *Lactobacillus* at species-level resolution, and in the context of the neighboring vaginal niche.

Our findings are also limited by several factors. Firstly, we used a cross-sectional design. Longitudinal sampling in the same participants before and after menopause, as well as before and after starting vaginal estrogen would have given a more comprehensive picture of the effects of menopause and hormones on urinary microbiota. While we did not perform these extended longitudinal studies, our findings are consistent with others who have shown differences in core urinary microbial taxa before and after menopause^31^. Another limitation is the lack of standardization of vaginal estrogen exposure. Though we required at least 6 weeks of therapy prior to sampling, participants may have been using vaginal estrogen for longer durations which could further influence *Lactobacillus* abundance data. A total of three participants in the Post+VE group were also using systemic forms of estrogen therapy (2 oral and 1 transdermal patch). While some investigators have concluded that systemic estrogens do not affect urinary conditions in the same manner as locally applied vaginal estrogen^39^, the effects of systemic estrogen on the urinary microbiome are debatable^40^ and thus could have led to skewed results in these few participants.

While our findings verified our hypotheses related to urinary lactobacilli, menopause, and vaginal estrogen, there are still many lingering questions. We identified several instances in which lactobacilli are found in high abundance in postmenopausal women not using estrogen. Specifically, in 9/61 Post participants who were not using estrogen therapy, *Lactobacillus* occupied >50% of the urinary microbiome. Most of these women also demonstrated high abundances of *Lactobacillus* in their vaginal microbiome. There are many questions about additional factors that lead to the presence of lactobacilli within the urinary tract. For example, we identified a strong inverse correlation between vaginal pH and *Lactobacillus* abundance in the bladder. Prototypically, a well estrogenized vaginal environment results in glycogen production by vaginal epithelial cells, a lower, more acidic pH, and subsequently the presence of vaginal lactobacilli^41^. Presumably, and supported by the paired data in our study, high abundances of vaginal lactobacilli would lead to higher abundances of the same species of *Lactobacillus* in the urinary bladder. Thus, there is a conceptual model supporting lowering vaginal pH to achieve downstream effects on lactobacilli within the urinary bladder. There are other agents used in clinical environments, such as boric acid, that may be able to achieve these effects without hormone^42^. In our study population there were several instances of acidic pH in postmenopausal women without estrogen, which was surprising. There are also several instances of premenopausal women with higher pH, which is less surprising given that sexual activity or other shifts in vaginal flora may contribute to a higher pH. However, understanding the various drivers for these differences in vaginal pH and how this indirectly influences urinary lactobacilli may ultimately help us to determine ways to optimize persistence of healthier bacterial populations within the bladder microbiome.

Lactobacilli exhibit properties that reduce pathogenic bacteria through several antimicrobial mechanisms. These include the production of lactate, other organic acids, bacteriocins, and surface-active molecules that promote aggregation into a “protective” biofilm^43,44^. *L. crispatus* is one species that has particularly been associated with urogenital (e.g., vaginal and urinary tract) health^43,45,46^. However, several species of *Lactobacillus* can counteract urinary pathogens *in vitro*^47^, and within the urinary bladder our findings support the premise that several species of *Lactobacillus* are associated with urinary tract health. It is unclear if lactobacilli exert a protective effect on the urinary tract simply through niche dominance, or whether there are species-dependent mechanisms that directly counteract pathogens. Metabolites are small biomolecules that play a key role in mediating host– microbe interactions. Studying metabolites may offer functional insight beyond microbial composition alone and can help to identify biochemical activities that distinguish closely related bacterial species. Our network correlational analysis of metabolites supports the assertion that *L. iners* behave differently than other lactobacilli within the urinary milieu, which mirrors findings from the vaginal environment^48^. Given that the majority of metabolites we identified were unannotated, it is not yet clear whether our findings reflect environments that support different *Lactobacillus* species or whether different *Lactobacillus* species are producing different metabolites. Future detailed studies elucidating functional implications of *Lactobacillus* species within the urinary microenvironment may be especially helpful.

## Conclusion

The urinary bladder microbial composition of postmenopausal women is significantly different than premenopausal women or postmenopausal women using vaginal estrogen, even when controlling for other potentially confounding variables. These differences are mainly driven by *Lactobacillus*, which is the most abundant genus in the urinary microbiome of Pre and Post+VE women. Urinary lactobacilli are positively correlated with vaginal lactobacilli, and often the same species of *Lactobacillus* are identified when testing paired bladder and vaginal samples from the same participants. More data are needed on which *Lactobacillus* species confer the greatest protection from pathogens, which key metabolic nutrients support these species, and how to best shape urinary microbial compositions to promote urologic health.

## Methods

### Participant selection and enrollment

This is a multi-institution cross-sectional analysis conducted under approval from Duke University Institutional Review Board (Pro00083917) and Oregon Health & Science University (OHSU) Institutional Review Board (IRB 10729). The analysis represents one of several planned analyses from the UMICRO study cohort, which enrolled 225 female participants (all self-identifying as women), including many with recurrent urinary tract infection (UTI). For this analysis, only those *<u>without</u>* recurrent UTI (see definition below) were considered, which comprised a subset of n=166 participants. **Supplemental Figure 1** shows the flow of participants from enrollment to analysis. Participants were recruited from urogynecology and urology clinics between August 2015 and March 2020.

### Inclusion and exclusion criteria

English speaking females who were premenopausal ages 30-48 (Pre), postmenopausal age > 55 years (Post), and postmenopausal age >55 years using vaginal estrogen for at least 6 weeks (Post+vagE2) were included. Exclusion criteria are detailed in **Supplemental Figure 1**; for this analysis we also excluded those with recurrent UTI during the 24 months prior to specimen collection. UTI was defined as urinary symptoms (e.g., dysuria, suprapubic pain, urinary frequency, or hematuria) accompanying a urine culture showing ≥ 10^4^ colony forming units (CFU) of a dominant organism. Recurrent UTI was defined as >3 UTI within 12 months or >2 UTI within 6 months^49^.

### Sample acquisition and clinical data collection

After informed consent, participants underwent transurethral catheterization and vaginal swab (distal vagina, just inside introitus) sampling per protocol, prior to any vaginal examination. Specimens were collected by trained personnel at a research visit. Vaginal pH was assessed from the same area where sampling occurred using pH paper. Demographic and medical history were collected, and participants completed questionnaires regarding sexual activity, urinary symptoms, dietary behaviors, medications, and supplements. Several clinical variables were collapsed into composite variables for analysis, as outlined in **Supplemental Table 2**.

### Clinical laboratory testing

Given that the original study cohort also included women with recurrent UTI (excluded for the current analysis), all participants underwent screening of urinary symptoms and laboratory testing at the time of sample collection. Chemical urinalysis was conducted. In the absence of clinical symptoms, specimens with nitrites, pyuria, or hematuria on chemical urinalysis were also sent for standard urine culture. Research study sample collection was only conducted in the absence of acute symptomatic UTI (based on symptoms and urinalysis). Those with asymptomatic bacteriuria identified by subsequent urine culture were excluded from downstream analyses.

### Specimen processing, DNA isolation, and sequencing

After chemical urinalysis, remaining urine was poured into 50 mL conical tubes. At OHSU, where the clinic is adjacent to the laboratory, whole urine was immediately transferred and frozen at −80°C, which was ultimately transferred to Duke. At Duke, where there are several clinics located remote from the laboratory, 50mL conical tubes contained DNA protectant (Assay Assure^TM^, Sierra Molecular Corporation, Incline Village, Nevada, USA) and were maintained at 4°C in the clinic until transfer to the laboratory within 24 hours. All urine samples underwent the same centrifugation and processing to separate the cellular fraction into pellets in one central laboratory, as previously described^13^. Vaginal swabs were maintained frozen at −80°C, and ultimately thawed, centrifuged, with cell pellets resuspended in enzymatic lysis buffer, per standard protocol instructions for Pretreatment for Gram-Positive Bacteria from the Qiagen DNeasy Blood and Tissue Kit (Qiagen, Valencia, California, USA), which was used with the lysozyme step for all DNA isolation. Positive [mock microbial community (Zymo Research, Orange, California, USA)] and negative (microbe-free water) controls were prepared in parallel with participant samples. DNA concentrations were measured using Qubit 4^TM^ Fluorometer with dsDNA HS Assay Kit^TM^ for Qubit (Thermo Fisher Scientific, Wilmington, Delaware, USA). Samples were submitted to the Duke Microbiome Shared Resource for library preparation and 16S rRNA amplicon sequencing via polymerase chain reaction (PCR) amplification of the V4 hypervariable region. To confirm specieslevel assignments for *Lactobacilli*, DNA including that from positive and negative controls were submitted to Loop Genomics (now Element Biosciences, Irvine, California, USA) for synthetic long read sequencing by LoopSeq^37^

### 16S rRNA amplicon data analysis

Sequencing data were bioinformatically processed using the DADA2 pipeline to identify taxa at a genus and species level. Briefly, DADA2 (v1.16.0) was used to infer amplicon sequence variants (ASVs) using default parameters^50^. BLCA (v2.2)^51^ was used with the NCBI 16S database (downloaded on 2021-11-05) to assign taxonomy to the species level. Samples that did not provide at least 2,000 reads per sample from sequencing data were excluded from further analysis, leaving analyzable sequencing data from the urinary bladder and vagina in n=149 and n=136 participants, respectively. The phylogenetic tree was built using Decipher (v 2.22.0) and phangorn (v 2.7.1). Decontam (v. 1.10.0) was evaluated for removal of contaminant DNA sequences, with a threshold *thr* of 0.5 used for the urine dataset. Sequencing data were rarefied to 2,000 for alpha and beta diversity analyses. Relative abundances of recovered taxa were visualized using the microshades package in R^52^

### Synthetic long read 16S rRNA data

Full-length contigs provided by Loop Genomics were processed using a DADA2 (v 1.26.0) pipeline adjusted for synthetic long read sequencing. Taxonomy was assigned in the same fashion as short read data, using BLCA (v2.2)^51^ with the NCBI 16S database (downloaded on 2021-11-05). Relative abundances were calculated for all bacteria identified and only bacteria that were assigned to a *Lactobacillus* genera were retained for further analysis (e.g., *Lactobacillus, Lacticaseibacillus, Limosilactobacillus*, etc. as defined in the new taxonomy presented in Zheng *et al*.^53^). Data were visualized using the microshades package in R^52^.

### Statistical Analysis

Microbial composition was compared between phenotypes using weighted UniFrac distance. Mean and median relative abundances were calculated for the top 10 genera from each niche (e.g., urinary bladder and vagina). Relative abundances of clinically relevant genera were compared by exposure group (e.g., Pre, Post, Post+VE) using Kruskal-Wallis with post hoc falsediscovery-rate (FDR) corrected pairwise testing using Wilcoxon rank sum. Pearson correlation coefficients were calculated to summarize correlations between relative abundances of *Lactobacillus* and age or vaginal pH. Four commonly used alpha diversity indices were calculated using the vegan^54^, phyloseq^55^, and microbiome^56^ R packages, including a measure of richness (number of observed species), two measures comprised of richness and evenness (the Shannon index and Inverse Simpson index) and a measure of evenness (Pielou’s evenness). Alpha diversity indices were compared between Pre, Post, and Post+VE groups using the Kruskal Wallis test followed by post-hoc pair-wise Wilcoxon Rank Sum tests with FDR correction. Beta diversity was evaluated with non-metric multidimensional scaling (NMDS) plots constructed from weighted UniFrac distance matrices to visualize community composition at the species level. Permutational multivariate analysis of variance (PERMANOVA) with the vegan adonis2 function and PERMANOVA R packages was used to compare UniFrac estimations while controlling for several covariates, as outlined in **Supplemental Table 2**.

### Urinary Metabolomics

During urine sample processing, urine supernatant was saved while pelleting the cellular fraction for DNA isolation. Banked urine supernatant was stored at −80°C, transferred to the University of California at San Diego (UCSD), and underwent untargeted metabolite analysis. Urine was combined with 100% MeOH for a final concentration of 80% MeOH. Samples were vortexed, incubated at −20°C for 20 min to allow for protein precipitation, then centrifuged for 15 min at 14,000 rpm before 800 μL of supernatant was transferred to a 96-Well DeepWell plate. Plates were dried and concentrated using a CentriVap Benchtop Vacuum Concentrator, covered and stored at −80°C until ready for LC-MS/MS data acquisition. For data acquisition, plates were resuspended in 200 μL of a 50% MeOH to water solution containing 1 μM of sulfadimethoxine as an internal standard. Untargeted metabolomics data were collected using an ultra-high-performance liquid chromatography system (Vanquish, Thermo Fisher Scientific, Waltham, MA, USA) coupled to an orbitrap mass spectrometer (Q-Exactive Hybrid Quadrupole-Orbitrap, Thermo Fisher Scientific, USA). The mobile phase solvent A consisted of 100% Optima LC-MS grade water with 0.1% formic acid, and the mobile phase solvent B consisted of 100% Optima LC-MS grade acetonitrile with 0.1% formic acid (LC–MS grade solvents, Fisher Chemical, USA). The flow rate was set to 0.500 mL/min with a 10 min per sample run time. The following gradient was used: 0 to 1 min held constant at 5% B; 1 to 7 min linear increase to 99% B; 7 to 8 min held constant at 99% B; 8 to 9 min linear decrease to 5% B; 9 to 10 min held constant at starting conditions. MS/MS data were collected with positive electrospray ionization mode.

### Metabolomic Analysis

Data in the .raw format were converted to .mzML files using the ProteoWizard tool MSConvert. All data (.raw and.mzML) were uploaded to MassIVE, saved under the ID MSV000095726 (ftp://massive.ucsd.edu/MSV000095726/). Feature finding was conducted using the Mzmine3 workflow version 3.9.2. MZmine feature table and .mgf outputs, along with sample metadata, were then input into the Global Natural Products Social Molecular Networking 2 (GNPS2) feature-based molecular networking (FBMN) tool for molecular networking and putative annotation through spectral library matching^57–59^. Finally, .graphml files were downloaded from the GNPS2 FBMN jobs and input into the free Cytoscape^58–60^ version 3.10.0 software to visualize molecular networks. Data were rclr-transformed using the vegan package version 2.6.10. Principal component analysis (PCA) and partial least squares discriminant analysis (PLS-DA) were performed with mixOmics version 6.26.0. Separation of group centroids in PCA space was tested using permutational multivariate analysis of variance (PERMANOVA). PLS-DA model performance was evaluated by 4-fold crossvalidation with 100 repeats. Ordination plots for both PCA and PLS-DA were generated using ggplot2 version 3.5.1. Variable importance in projection (VIP) scores were extracted from the PLS-DA model, and features with VIP > 1 that also carried FBMN annotations were retained for downstream analysis.

### Multi-omics Analysis

Data preparation and reformatting were performed using the FBMNStats application^38^. Feature tables were filtered to retain only the samples belonging to each experimental group, followed by blank removal and imputation of remaining missing values. Three separate feature tables were generated for each experimental group, then each table was then analyzed independently. Multi-omics integration of microbial and metabolomic data was performed using CorrOmics. For each submission, metabolomic data were filtered by the “ATTRIBUTE_Corromics_filename” attribute, log10-transformed with 0-to-1 imputation of missing values, and subjected to correlation analysis using the Pearson method. Correlation edges were then filtered using a target-decoy false discovery rate (FDR) approach, retaining only correlations stronger than ±0.50. Filtered correlation networks from CorrOmics were imported into Cytoscape where visual properties were mapped to feature attributes to aid interpretation. VIP scores and GNPS-FBMN annotations were imported into Cytoscape as separate node attribute tables and matched to the network nodes by shared feature ID. Node size and color indicated annotation status while node border color indicated whether the metabolite had a VIP score > 1. Edge width was mapped to the strength of the correlation, and edge color was used to distinguish positive from negative metabolite-microbe correlations. Metabolites that were correlated but lacked either a VIP score > 1 or annotation were removed from the network unless correlated with more than one microbial species.

## Acknowledgments

We thank the study participants for their invaluable contributions to this research and the clinical teams for their assistance with participant recruitment and specimen collection. We are grateful to Zhuoqun (Carol) Wang and Alex Rouhier for their contributions to study support and project activities. We thank Silvia Grant-Beurmann, Ph.D., for her review and editing of the manuscript.

This work was supported by the National Institute of Diabetes and Digestive and Kidney Diseases through K01DK116706 (L.K.), K23DK110417 (N.Y.S.), and by the National Institute on Aging through R03AG060082 (N.Y.S.). These awards supported the development of the study, generation of preliminary data, and completion of the work presented in this manuscript.

## Notes

### Competing Interest Statement

NYS has received research support from Medtronic Inc, Ethicon J&J, Locus Biosciences, and Caldera Medical, as well as royalties from UpToDate.

