## Supplemental Data for "The Urogenital *Lactobacillus* Axis Across Menopause: Connections Between the Vagina, Bladder, and Urinary Metabolome"

**Supplemental Table 1:** V4 16S rRNA sequencing read depth and taxonomy by niche

| **Metric** | | **Urinary Bladder Samples** | **Vaginal Samples** |
| --- | --- | --- | --- |
| Median read depth | | 32,067 | 45,320 |
| Minimum read depth | | 2,054 | 19,444 |
| Maximum read depth | | 91,168 | 92,652 |
| # Identified Taxa | |  |  |
|  | Phyla | 16 | 12 |
|  | Classes | 42 | 33 |
|  | Orders | 91 | 61 |
|  | Families | 208 | 126 |
|  | Genera | 500 | 301 |
|  | Species | 895 | 526 |

**Supplemental Table 2: Clinical Covariates Used in PERMANOVA models**

| **Variable** | **Individual Variables Collected for Composite Variables** | **Notes** |
| --- | --- | --- |
| Daily probiotics/yogurt | Yogurt (regular basis) Probiotics daily | Included if participant indicated “yes” to either |
| Race/ethnicity | Race collected as Caucasian, African American, Asian, Native American, or Multi-racial. Ethnicity collected as Latina or non-Latina. | Categorical data described in Table 1; collapsed into two categories (white & non-white) for statistical models. |
| Sexually Active |  | Participant self-report |
| OAB (overactive bladder) | Composite of *either* medical diagnosis (urgency urinary incontinence or OAB) from chart review *or* known OAB medication use (e.g., bladder anti-muscarinic or beta agonist oral medications) | Did not specify “wet” (e.g., urgency urinary incontinence) or “dry” OAB |
| BMI (body mass index) | Calculated from weight and height as kilograms/meters^2^ | Collected from medical record |
| Diabetes |  | Participant self-report on comorbidity questionnaire or medical diagnosis attributed in medical record |
| Age |  | In years, on date of specimen acquisition |

**Supplemental Table 3: Top Genera Recovered in Urinary Bladder & Paired Vaginal Samples**

|  | **Premenopausal**  **N=28** | | |  | **Postmenopausal +Vaginal Estrogen**  **N= 36** | | |  | **Postmenopausal**  **N = 60** | | |
| --- | --- | --- | --- | --- | --- | --- | --- | --- | --- | --- | --- |
|  | **Genus** | **n present (%)** | **Median Relative Abundance** |  | **Genus** | **n present (%)** | **Median Relative Abundance** |  | **Genus** | **n present (%)** | **Median Relative Abundance** |
| **URINARY BLADDER** | Lactobacillus | 27 (96) | 42.93 |  | Lactobacillus | 30 (83) | 7.53 |  | Vibrio | 55 (92) | 5.98 |
|  | Vibrio | 24 (86) | 2.53 |  | Vibrio | 34 (94) | 5.73 |  | Halomonas | 56 (93) | 4.98 |
|  | Halomonas | 25 (89) | 1.93 |  | Halomonas | 34 (94) | 3.43 |  | Stenotrophomonas | 51 (85) | 2.70 |
|  | Gardnerella | 20 (71) | 0.80 |  | Pseudomonas | 33 (92) | 1.70 |  | Pseudomonas | 51 (85) | 2.53 |
|  | Klebsiella | 25 (89) | 0.68 |  | Klebsiella | 34 (94) | 1.68 |  | Klebsiella | 52 (87) | 2.08 |
|  | Stenotrophomonas | 20 (71) | 0.50 |  | Stenotrophomonas | 33 (92) | 1.13 |  | Lactobacillus | 42 (70) | 0.55 |
|  | Pseudomonas | 18 (64) | 0.30 |  | Enterococcus | 31 (86) | 0.40 |  | Escherichia | 46 (77) | 0.48 |
|  | Corynebacterium | 18 (64) | 0.10 |  | Pantoea | 28 (78) | 0.28 |  | Enterococcus | 47 (78) | 0.45 |
|  | Escherichia | 18 (64) | 0.10 |  | Brucella | 27 (75) | 0.18 |  | Streptococcus | 47 (78) | 0.38 |
|  | Enterococcus | 15 (54) | 0.05 |  | Alishewanella | 22 (61) | 0.12 |  | Brucella | 41 (68) | 0.30 |
| **VAGINA** | Lactobacillus | 27 (96) | 88.3 |  | Lactobacillus | 32 (89) | 73.1 |  | Lactobacillus | 51 (85) | 2.0 |
|  | Gardnerella | 16 (57) | 0.8 |  |  |  |  |  | Streptococcus | 44 (73) | 1.3 |
|  |  |  |  |  |  |  |  |  | Anaerococcus | 43 (72) | 0.6 |
|  |  |  |  |  |  |  |  |  | Finegoldia | 43 (72) | 0.3 |
|  |  |  |  |  |  |  |  |  | Corynebacterium | 38 (63) | 0.2 |
|  |  |  |  |  |  |  |  |  | Dialister | 36 (60) | 0.1 |
|  |  |  |  |  |  |  |  |  | Halomonas | 31 (52) | 0.1 |
|  |  |  |  |  |  |  |  |  | Vibrio | 33 (55) | 0.1 |

Table 3 shows the top 10 genera identified in paired urinary and vaginal samples from n = 124 participants. Genera from the urinary bladder are listed in order of median relative abundance per study group. Since each genus is not be found in every sample, the proportion of samples where a particular genus was recovered is noted. The top 10 genera identified in the vagina are also noted. Given substantial skew & sparsity in microbial data, median abundances are 0 in many instances, and only those >0 are listed.

**Supplemental Table 4: *Lactobacilli* recovered with synthetic long read sequencing technology**

| *Lactobacillus iners* | *Lactobacillus crispatus* | *Lactobacillus gasseri* |
| --- | --- | --- |
| *Lactobacillus jensenii* | *Lacticaseibacillus casei* | *Lactobacillus hominis* |
| *Lactobacillus taiwanensis* | *Lactobacillus kefiranofaciens* | *Lacticaseibacillus paracasei* |
| *Lactobacillus colini* | *Lactobacillus gallinarum* | *Limosilactobacillus frumenti* |
| *Limosilactobacillus pontis* | *Lactobacillus fornicalis* | *Lactobacillus ultunensis* |
| *Lactobacillus delbrueckii* | *Lactobacillus johnsonii* | *Limosilactobacillus fermentum* |
| *Lactobacillus helveticus* | *Limosilactobacillus vaginalis* | *Limosilactobacillus coleohominis* |
| *Lacticaseibacillus rhamnosus* | *Limosilactobacillus reuteri* | *Lactobacillus psittaci* |
| *Lactobacillus hamsteri* | *Lactobacillus acidophilus* | *Limosilactobacillus antri* |
| *Lactobacillus equicursoris* | *Lactobacillus gigeriorum* | *Lactobacillus amylovorus* |
| *Lactobacillus kalixensis* | *Lactobacillus amylolyticus* | *Lactobacillus pasteurii* |
| *Levilactobacillus hammesii* | *Lactiplantibacillus plantarum* | *Lactobacillus intestinalis* |
| *Lactobacillus acetotolerans* | *Amylolactobacillus amylotrophicus* | *Lactobacillus rodentium* |
| *Lactobacillus helsingborgensis* | *Secundilactobacillus kimchicus* | *Limosilactobacillus caviae* |
| *Lactiplantibacillus paraplantarum* | *Liquorilactobacillus cacaonum* | *Limosilactobacillus ingluviei* |
| *Lactobacillus bombicola* | *Lactobacillus kitasatonis* |  |

Detailed *Lactobacillus* species identified in n=71 urinary samples re-sequenced using a synthetic long-read approach that covers the entire 16S rRNA gene. What was once termed as *Lactobacillus* are now 24 different genera including *Limosilactobacillus*, *Lacticaseibacillus*, and several others.^51^

**Supplemental Figure 1: Recruitment Flow Diagram**

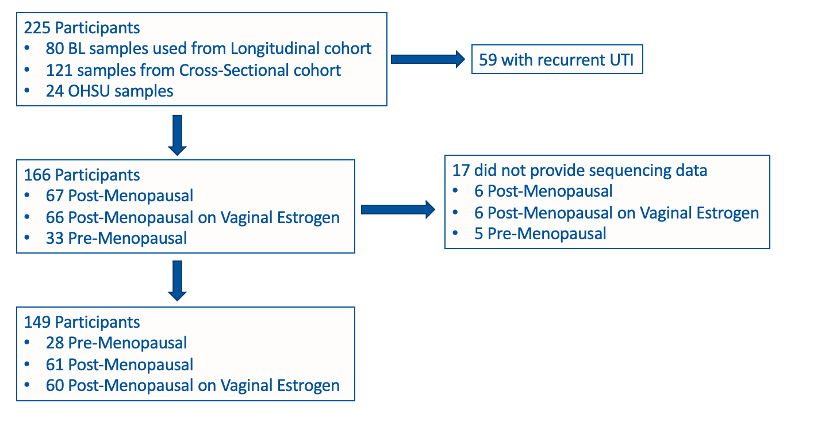

*

*Participants compiled from UMICRO study (Duke University) which had a longitudinal cohort of postmenopausal women with recurrent UTI and matched controls (baseline samples from non-recurrent UTI participants included here), and a cross-sectional cohort sampled only for this analysis. A separate cross-sectional cohort from Oregon Health & Science University (OHSU) were sampled using the same protocol. Exclusion criteria (for Longitudinal and Cross-Sectional cohorts) included: 1) those who did not fall into the required age groups (e.g., younger than 30 or between 48-54 years); 2) instrumentation of the urinary tract (e.g., cystoscopy) within the prior 30 days; 3) neurogenic bladder dysfunction or any need for chronic catheterization; 4) pregnant or breastfeeding within the prior 6 months; 5) intravaginal pessary use; 6) renal insufficiency (creatinine >1.3); 7) diagnosis of interstitial cystitis/painful bladder syndrome; 8) active malignancy; 9) history of prior pelvic radiation; 10) breast cancer diagnosis within the prior 5 years or on any anti-estrogen therapy

**Supplemental Figure 2: Paired urinary bladder & vaginal microbial data (all bacteria)**

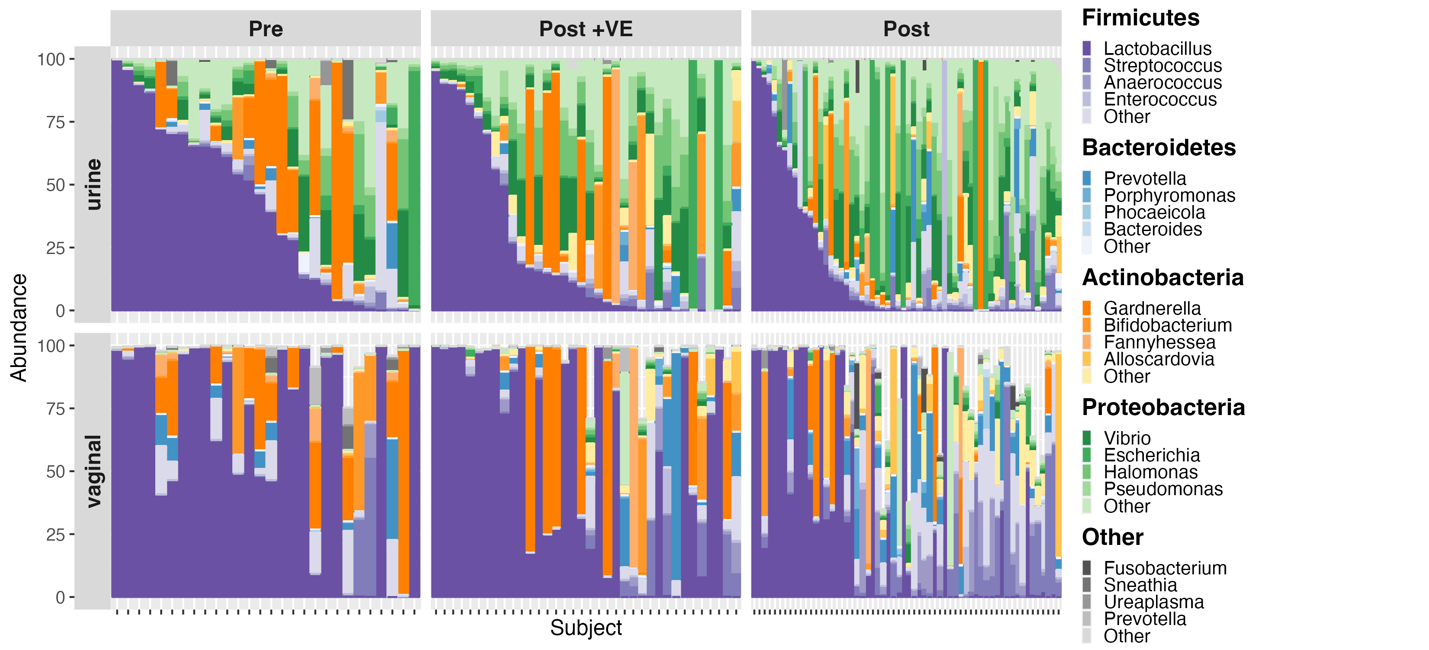

Stacked bar plots depicting recovered microbes from V4 16S rRNA amplicon sequencing at the at the genus level in n=124 participants with paired samples. Data are sorted in order of relative abundance of *Lactobacilli* in the urinary bladder and aligned showing microbiota from the urinary bladder (top) and vaginal (bottom) samples.

**Supplemental Figure 3: Paired urinary bladder & vaginal microbial data (Actinomycetia)**

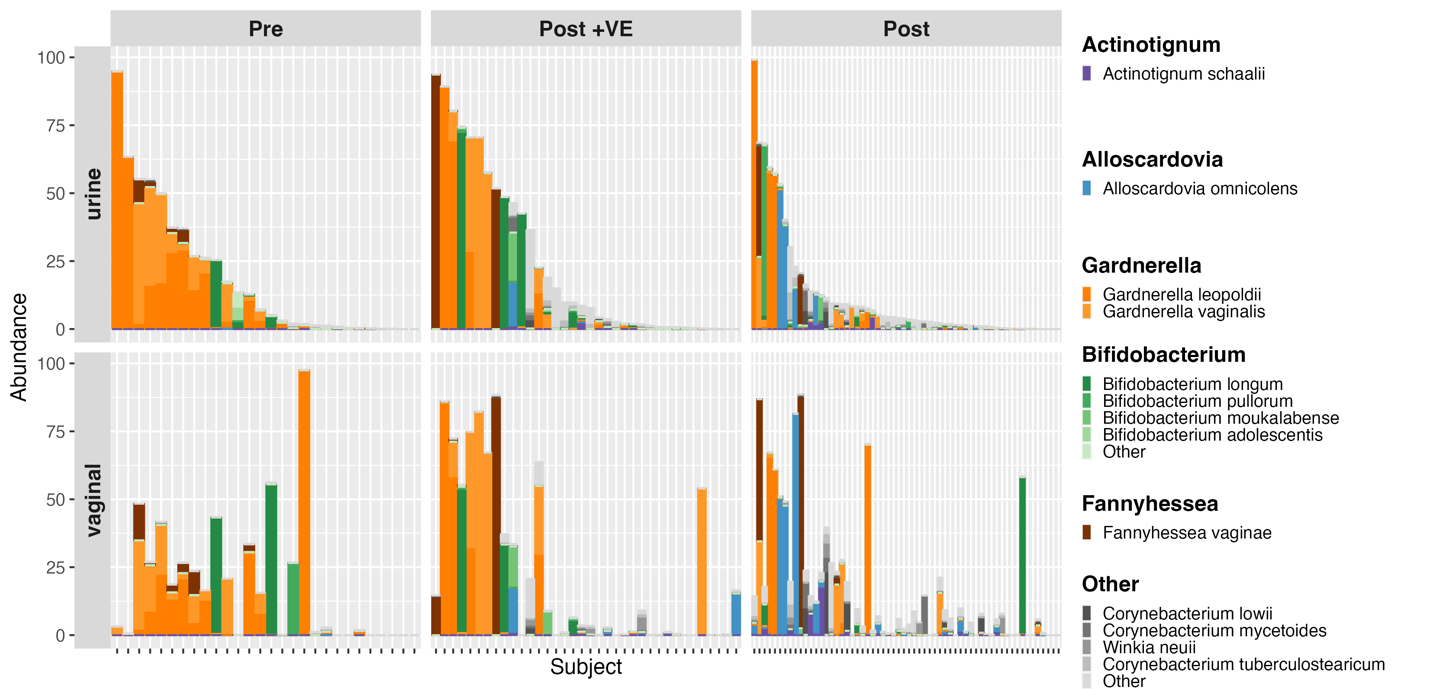

Figure depicts stacked bar plots from paired urinary bladder (top) and vaginal (bottom) samples from the same participants. Data are sorted by relative abundance of the bacterial class Actinomycetia within the urinary bladder, and aligned, with different genera represented by color as noted in the key. For genera from this taxonomic class, presence in one niche (e.g., urinary bladder) is correlated with the adjacent niche (e.g., vagina).

**Supplemental Figure 4: Paired urinary bladder & vaginal microbial data (Enterobacteriaceae)**

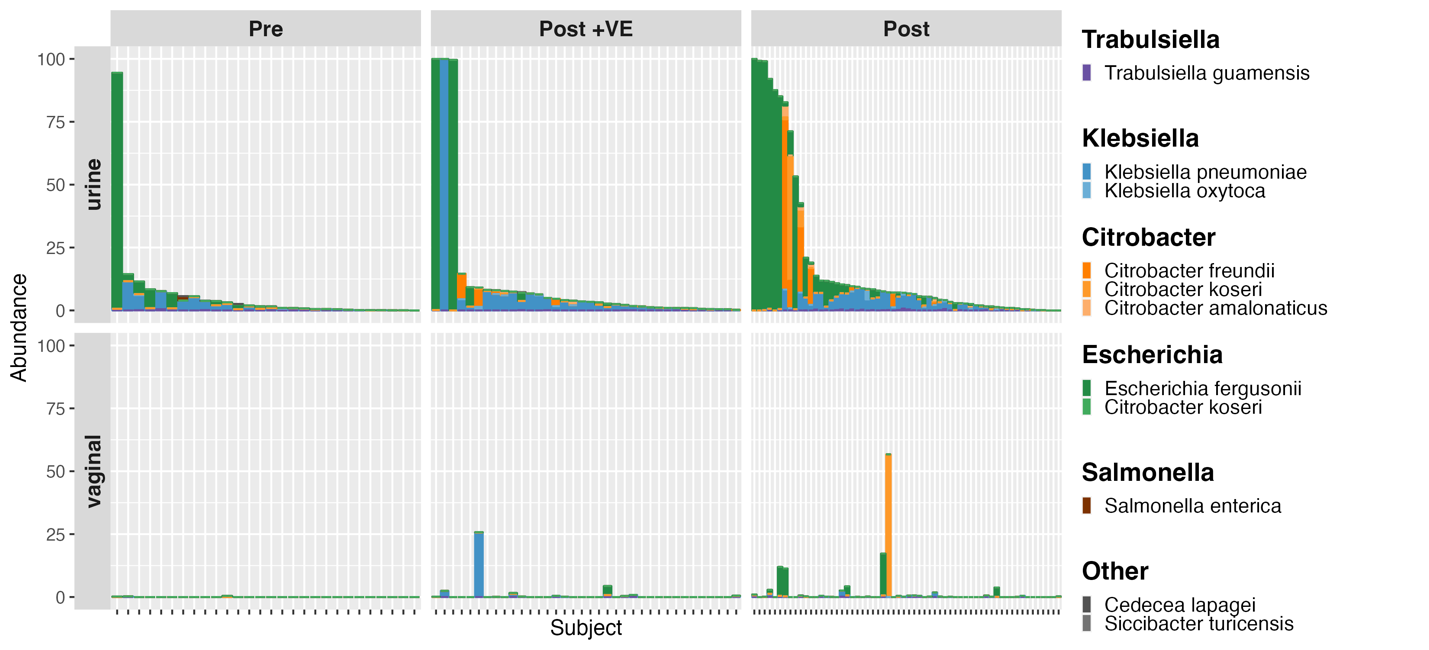

Figure depicts stacked bar plots from paired urinary bladder (top) and vaginal (bottom) samples from the same participants. Data are sorted by the relative abundance of the bacterial family Enterobacteriaceae within the urinary bladder, then aligned, with different genera represented as colors noted in the key. For this taxonomic family, correlations between presence in the urinary bladder and vagina are not present or are hard to visualize due to low abundances of these bacterial taxa overall in the vagina.
